# Ab initio derivation of the FRET equations resolves old puzzles and suggests measurement strategies

**DOI:** 10.1101/394635

## Abstract

Quantitative FRET-based imaging methods rely on the determination of an apparent FRET efficiency (*E*_*app*_) as well as donor and acceptor concentrations, in order to uncover the identity and relative abundance of the oligomeric (or quaternary) structures of associating macromolecules. Theoretical work has provided “upwards” relationships between the experimentally determined *E*_*app*_ distributions and the quaternary structure models that underlie them. By contrast, the body of work that predicates the “downwards” dependence of *E*_*app*_ on directly measurable quantities (i.e., fluorescence emission of donors and acceptors) relies largely on plausibility arguments, one of which is the seemingly obvious assumption that the fraction of fluorescent molecules in the ground state pretty nearly equals the total concentration of molecules. In this work, we use the kinetic models of fluorescence in the presence and absence of FRET to rigorously derive useful relationships between *E*_*app*_ and measurable fluorescence signals. Analysis of these relationships reveals a few anticipated surprises and some unexpected explanations for known experimental FRET puzzles, and it provides theoretical foundations for optimizing measurement strategies.

## I. INTRODUCTION

Förster Resonance Energy Transfer (FRET) (1-3) is, without doubt, a very useful physical phenomenon that enjoys broad popularity among researchers in various science areas (4-8). Defined as the transfer of energy from an excited fluorescent tag to an unexcited one, both of which are attached to macromolecules of interest *in vivo* or *in vitro*, FRET has become an indispensable tool in a gamut of applications ranging from estimation of intra-molecular distances within a protein or DNA molecules (2, 9), through probing the structure of oligomeric complexes (10-13) and to the determination of the proportion of various oligomeric species and their dissociation constants in living cells (11, 14). Fully quantitative analysis has been facilitated by the use of the kinetic theory of FRET (15, 16) as well as the availability of multiphoton microscopy with spatial and spectral resolution (7, 10, 12, 17, 18).

A relevant quantity in FRET is the efficiency of energy transfer (*E*), which depends on the sixth power of the ratio between the Förster distance, *R*_*0*_, and the distance between the chromophores of the fluorescent tags (1, 19, 20). Customarily, *E* is connected to the lifetimes of the excited state of the donor in the presence (τ^*DA*^) and absence (τ^*D*^) of acceptors in fluorescence lifetime imaging (FLIM) measurements, as well as to the fluorescence emitted by the donor in the presence (*F*^*DA*^) and absence (*F*^*D*^) of acceptors in steady-state intensity-based measurements (17, 20, 21).

In typical FRE T experiments, *F*^*DA*^ or τ^*DA*^ are measured separately from *F*^*D*^ or τ^*D*^, although preferably they should be measured from the same sample after FRET is somehow abolished, such as by inducing separation of the molecular complexes. The latter is rarely, if ever, possible, and several methods have been devised, which provide different degrees of approximation to the true values of *F*^*D*^ and τ^*D*^, and therefore of *E*. Such methods rely on various corrections (for example, for spectral bleed-through and acceptor photo-bleaching), as reviewed by Jares-Erijman and Jovin (22). Alternatively, one avoids use of corrections or additional measurements altogether by using spectral resolution to quantify a reduction in the donor emission as well as acceptor sensitized emission simultaneously, thereby separating the donor and acceptor signals upon a single sample scan (10, 23-25). This latter approach has led to the introduction of different variants of FRET-based imaging, including FRET spectrometry (12, 14, 26), fully quantitative spectral imaging (FSI) (11, 13) and, more recently and only theoretically for now, FRET-induced color contrast shift (FiCoS) spectrometry (27).

It is generally recognized that, when a single excitation wavelength is used, it is possible to determine the FRET efficiency and the donor concentration, but not the acceptor concentration (11, 12, 14, 27). Experimentally determined distributions of apparent FRET efficiencies (*E*_*app*_) comprise one or more peaks, each of which can be simulated using a certain proportion of donor and acceptor molecules within an oligomeric complex with a certain size and geometry (20, 28). The model that correctly predicts the number and position of each peak in the *E*_*app*_ histogram is taken as the quaternary structure of the protein of interest. This method is known as “*FRET spectrometry*.” When used in conjunction with a second excitation wavelength it also provides the acceptor concentration in addition to *E*_*app*_ and donor concentration (7, 11, 12, 14, 17), which could be used to monitor any dependence of the oligomer size or geometry on concentration (12, 14).

The relationships between the experimentally determined *E*_*app*_ and the theoretical models incorporating the geometry and size of the oligomer as well as the different possible proportions of donors and acceptors within each complex are termed in this paper “*upwards relationships*” and are based on the well-tested (16, 29) kinetic theory of FRET (15). By contrast, “*downwards relationships*” which allow one to compute *E*_*app*_ from donor and acceptor signals, without making arbitrary assumptions about the probabilities to find molecules in excited or ground states, have not been rigorously derived from the kinetic model of FRET until now. The work presented in this report starts from suitable kinetic models for fluorescent molecules in the presence and absence of FRET to derive expressions for *E*_*app*_ as well as donor and acceptor concentrations from fluorescence emission of acceptors and donors, some of which have been introduced and used but not rigorously derived in previous publications (10, 15, 20). It is shown that some of the assumptions that were implicit in some of the downwards relationships used by all of us in the FRET community, are not automatically valid and could lead to systematic errors if not carefully considered in the context of the experimental protocol used. Detailed analysis of those assumptions allows us to provide long-sought explanations for some known experimental puzzles. In addition, it provides the theoretical basis for identifying and refining the experimental conditions for quantitative FRET methods based on intensity measurements with no temporal resolution. A possibility is also suggested for using temporally resolved measurements for computing *E*_*app*_ distributions needed in FRET spectrometry analysis.

## II. THE KINETIC MODELS OF FLUORESCENCE AND FRET

### II.1. The kinetic equations

The relationships between different de-excitation rates and probabilities of fluorescent molecules to be in their excited states are obtained from the simple kinetic models presented in Figure 1. According to these models, for donors in the absence of acceptors (Figure 1a) the rate of change of the probability (*p*_*D*_*\**) to find a donor in an excited state (*D*^*\**^) is

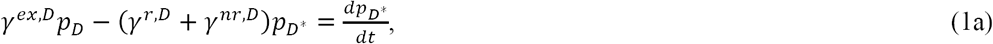

where *p*_*D*_ is the probability to find a donor in its ground state, *γ*^*ex,D*^ is the rate of excitation of donors initially in their ground state (*D*), while *γ*^*nr,D*^ and *γ*^*r,D*^ are the rates of donor de-excitation through non-radiative (e.g., internal conversion) and radiative (i.e., emission of a photon) processes, respectively. Similarly, for acceptors in the absence of donors (Figure 1b) the rate of change of the probability to find the *j*-th acceptor in an excited state (*p*_*A*,j*_) is

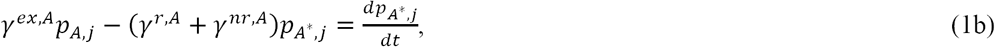

where *p*_*A,j*_ is the probability to find the *j*-th acceptor in its ground state, *γ*^*ex,A*^ is the rate of excitation of acceptors in their ground state (*A*), while *γ*^*nr,A*^ and *γ*^*r,A*^ are the rates of acceptor de-excitation through non-radiative and radiative processes, respectively.

**Figure 1.**
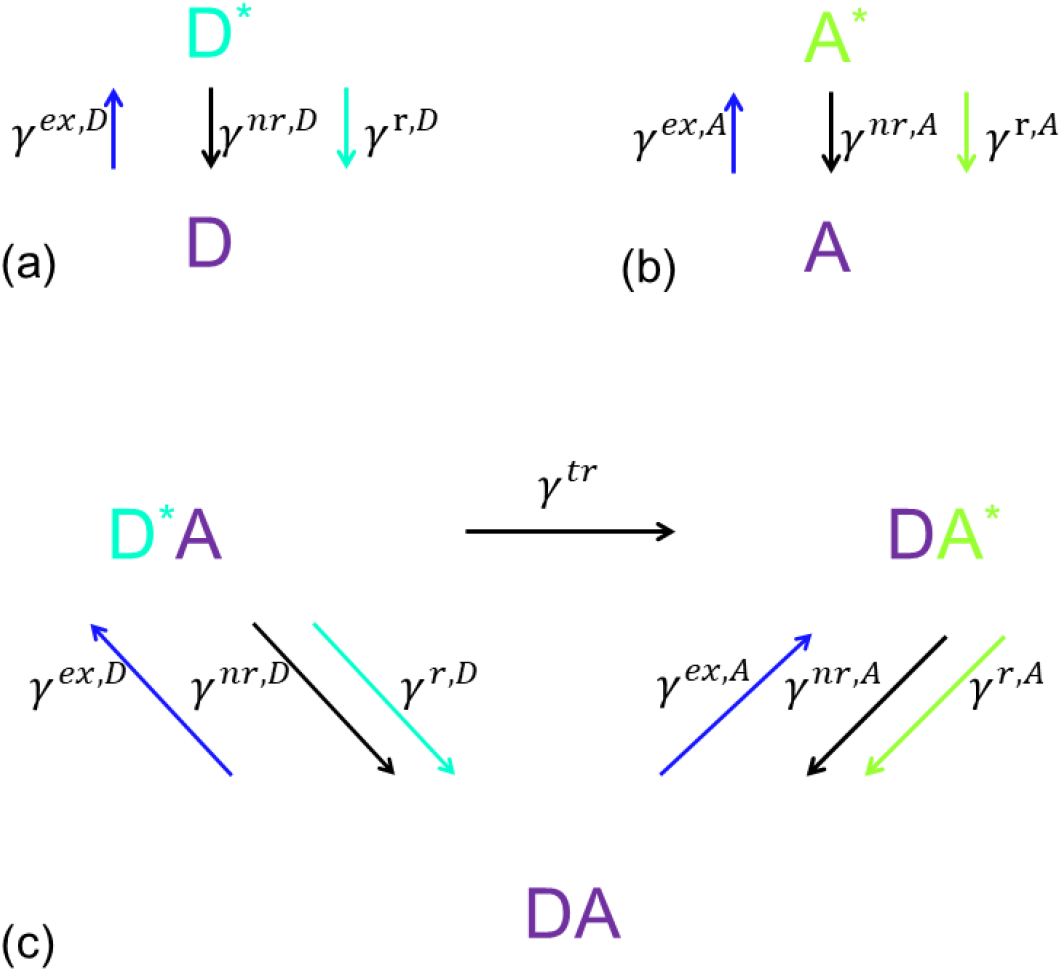
Kinetic models of fluorescence and FRET for (a) donors, D, in the absence of acceptors, A, (b) acceptors in the presence of donors, as well as (c) donors and acceptors in the presence of each other (i.e., FRET). Asterisk denotes excited species. All other symbols are defined in the text.

For single donors associated with *n* acceptors, if the acceptors are excited both directly by light and through energy transfer from the donor, the kinetic model of FRET (Figure 1c) gives:

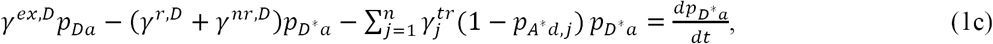

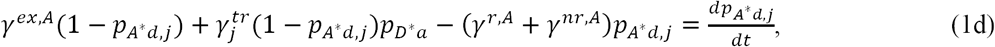

where 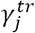 is the rate of energy transfer to each of the *n* acceptors, while *p*_*Da*_, *p*_*D*a*_, *p*_*Ad,j*_, and *p*_*A*d,j*_ are the probabilities to find donors and acceptors in their unexcited or excited states in the presence of each other. These probabilities obey the relations: *p*_*Da*_ + *p*_*D*a*_ = 1 and *p*_*Ad,j*_ + *p*_*A*d,j*_ = 1.

For single photon excitation, the excitation rates of donors and acceptors depend on the light irradiance, I (in W/m^2^), and the wavelength-dependent extinction coefficient of each fluorescent species, ε^*X*^(λ_*ex*_), according to the expression

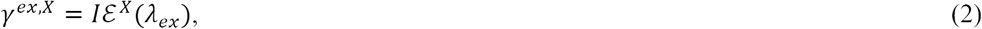

where *X* stands for *A* or *D,* and *ε*^*X*^ = ε^*X*^(λ_*ex*_)In(10)λ_*ex*_*(hcN*_*A*_)^−1^, with λ_*ex*_ being the excitation wavelength, *h* Plank’s constant, *c* the speed of light, and *N*_*A*_ Avogadro’s number. An instrumental multiplicative factor also may be incorporated, to account for changes introduced by the optics, but this almost invariably leads to a change in the excitation light intensity at the position of the sample, and so it can be safely absorbed into *I*, for simplicity.

As it is well known, for two-photon excitation, the extinction coefficient depends on the second power of the light intensity. We are not going to consider that particular fact herein, since from the point of view of this research, the effect of two-photon excitation is only to change the value of the excitation rate and not the kinetics behavior.

### II.2. Particular forms for long integration time and no temporal resolution

Taking Eq. (2) into account and integrating Eqs. (1) with respect to time, we have:

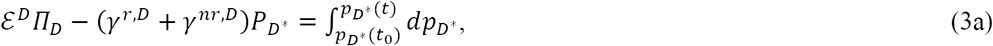

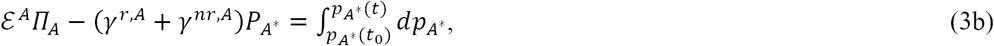

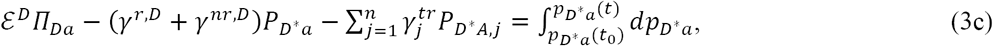

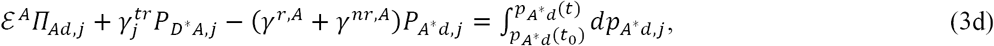

where we have used the following notations for the various integrals: 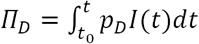, 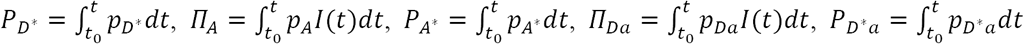, 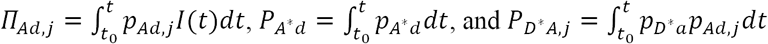.

The probability to find donors or acceptors in their excited states is always less than unity, while the excitation and de-excitation rates are all greater than 10^4^ s^−1^ (with de-excitation rates being some three to five orders of magnitude higher). As a result, for integration times of 1 μs or greater, the values of the integrals on the left-hand-side of Eqs. (3) far exceed the values of the integrals on the right-hand-side, which are always ≤ 1. This applies both to the case of steady-state intensity measurements employing continuous-wave light sources, where the probabilities are constant, and to measurements employing ultra-short light pulses that change the excited state probabilities from zero to some higher value and then back to zero. In either case, the integrals on the right hand side of Eqs. (3) vanish, and we have:

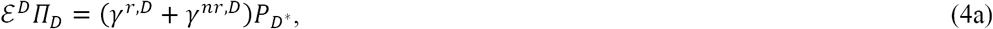

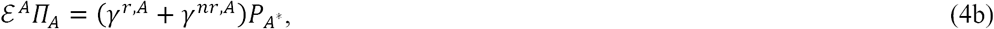

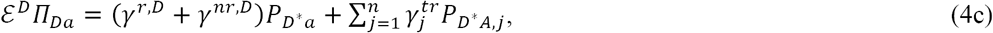

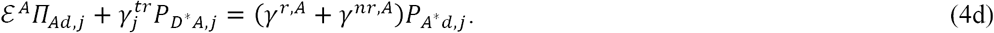

#### II.2.1 Continuous wave excitation and long integration time

If continuous wave (CW) lasers are used, the light intensity, *I*, is constant on timescales larger than those corresponding to the statistical fluctuations of the photons of light, and Eqs. (4) become

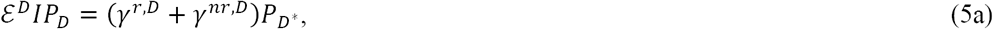

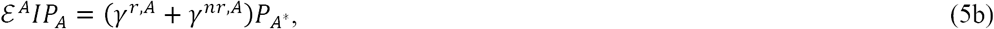

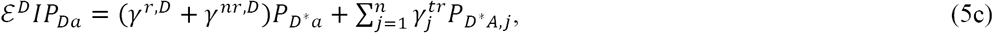

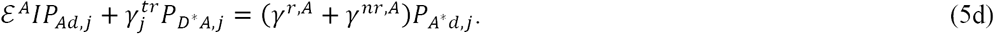

where the probabilities on the left-hand-side of these equations, 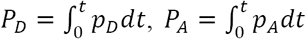, 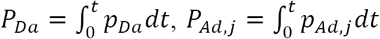, are now constant. These equations will be used later on in this paper to express the emission intensities of donors and acceptors as a function of rate constants and other experimental parameters for continuous wave excitation, i.e., steady state studies.

#### II.2.2 Pulsed excitation and long integration time

If the excitation consists of trains of light pulses that are much shorter than the excited lifetime of the fluorescent species, the intensity may be, to a good approximation, described by

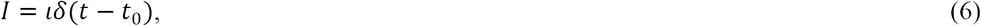

where *ι* is a dimensional coefficient, and *δ(t)* is Dirac’s delta function. Using this formula and the sifting property of the Dirac delta function^1^ in the П integrals defined above, Eqs. (4) become:

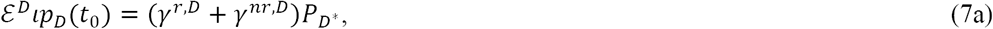

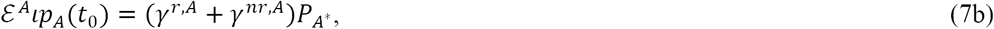

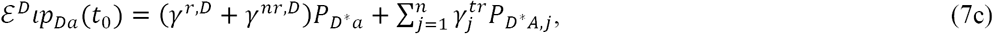

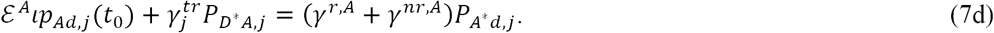

The interpretation of the assumption used in the derivation of Eqs. (7) is that the ultrashort pulse applied to a sample at the initial time *t*_0_ brings a fraction of the donors or acceptors into their excited states within a time much shorter than the lifetime of the donor, i.e., well before any de-excitation process starts. The excitation/de-excitation cycle repeats itself at the repetition rate of the laser light pulses for the entire duration of the data collection (or integration time). If each pulse arrives at the sample a long time after the previous one (relative to the fluorescence lifetime of the molecules), it finds all the molecules in their ground states. In this case, we may assume that *p*_*D*_*(t*_0_) = *p*_*Da*_*(t*_0_) = *p*_*A*_*(t*_0_) = *p*_*Ad*_*(t*_0_) = 1, and Eqs. (7) become:

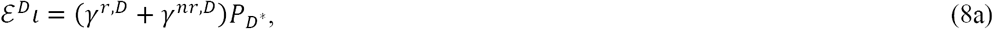

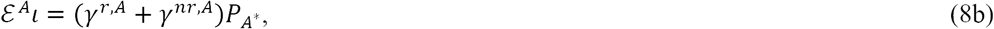

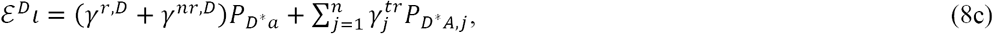

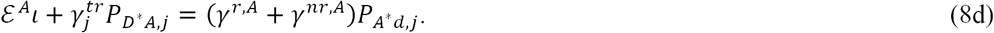

Equations (8) will be used later to express the integrated emission intensity of the differing fluorescent species in the case of pulsed excitation and acquisition times much longer than the repetition rate of the light pulses, i.e., in the absence of temporal resolution.

### II.3 Particular forms for pulsed excitation and temporal resolution

If an ultrashort pulse, which may be described by Eq. (6) is applied to the sample, and then the fluorescence emission is measured repeatedly for integration times much shorter than both the lifetime of the fluorescent molecules and the time elapsed between arrival of two successive pulses, fluorescence decay curves may be acquired for each fluorescent species, such as donors only or donors in the presence of acceptors. Typical values are: 10 to 100 ps for integration times, a few ns for the excited state lifetime, and ten or more ns for the repetition rate of the laser pulses.

#### II.3.1 Case A: acceptors are not excited by light

If the ultrashort pulse has a center wavelength that only matches the excitation maximum of the donor and does not excite the acceptor to any significant extent, then *p*_*Ad,j*_ = 1. Under these conditions, using Eq. (2) and separation of variables, Eqs. (1a) and (1c) become

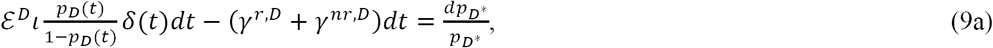

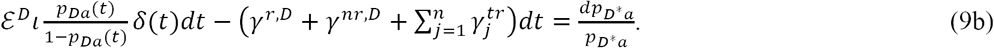

Upon integrating the left side of Eqs. (9) from 0 to *t* and the right side from *p_D*_*(0) [or *p*_*D*a*_(0)] to *p*_*D\**_(*t*), the sifting property of the Dirac delta function now provides that the first terms on the left-hand-sides vanish (since the lower limit of integration is not less than zero), and we therefore obtain:

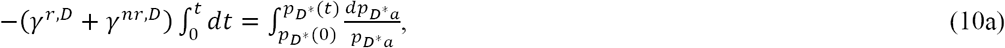

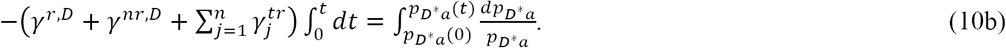

Since each ultrashort pulse brings a fraction of donors in their excited state well before any de-excitation occurs, we have *p*_*D\**_(*t*), *p*_*D*a*_(*t*) → *p*_*D\**0_ when *t* → 0 for all the donors regardless of whether there are any acceptors nearby. In addition, the probabilities that donors are in excited states tend to zero, i.e., *p*_*D\**_(*t*), *p*_*D*a*_*(t)* → 0, for very long times, *t* → ∞. This upper limit is only infinite by comparison to the donor lifetime and is in fact equal to the time elapsed between the arrival of successive light pulses. With these boundary conditions, Eqs. (10) admit the following solutions:

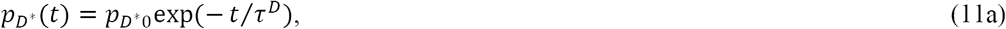

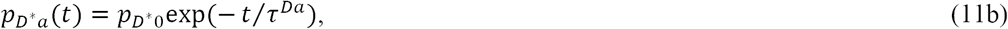

in which

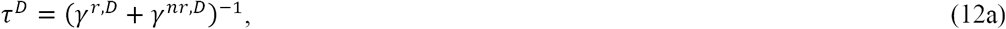

and

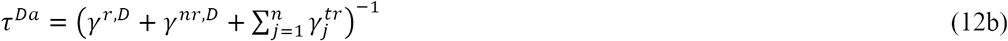

are the *donor lifetimes* in the absence and presence of acceptors, respectively, while

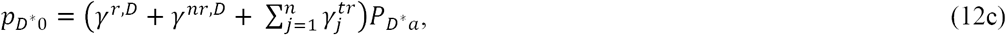

is the total number of excitations, which is equal to the sum of excitations lost through all de-excitation pathways. Eq. (12c) may be derived by integrating Eq. (1c) (assuming *p*_*Ad,j*_ =1) from 0 to a very long time, and the probability limits of *p*_*D\**0_ and 0.

#### II.3.2 Case B: acceptors are directly excited by light

When a light pulse described by Eq. (6) excites not only the donors but also the acceptors, competition occurs between energy transfer and laser light for exciting the acceptors, and hence *p*_*Ad,j*_ < 1. Under this condition and following the procedure described in Case A, integration of Eq. (1c) gives:

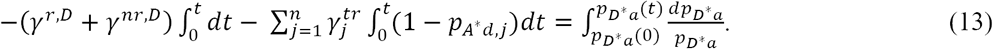

This type of equation has the following analytical solution for *p*_*A*d,j*_*(t)* functions that have the property *p*_*A*d,j*_ = *p*_*A\**0_φ_*A*d,j*_*(t)*:

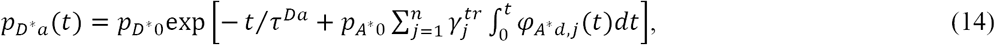

where *p*_*A\**0_ is an initial amplitude and φ_*A*d,j*_ is an arbitrary decay function (which would take the form of an exponential decay when there is only direct excitation of the acceptors). The integral in the exponent may be evaluated numerically from experimentally determined acceptor fluorescence decay, and

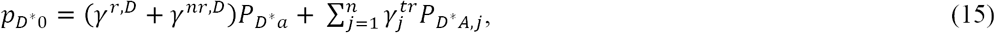

is the total number of donor excitations, expressed as the total number of excitations lost through different de-excitation pathways. Equation (15) may be derived rigorously by integrating equation (1c) over time from 0 to very long time, which corresponds to the probability limits of *p*_*D\**0_ and 0. Obviously, Eq. (14) reduces itself to Eq. (11b) when there is no direct excitation of the acceptor by light, i.e., for *p*_*A\**0_ = 0.

### II.4 Fluorescence emission in the presence and absence of FRET

#### II.4.1 The fundamental equations of FRET

The quantities of interest in FRET are the quantum yields of the donor in the absence (*Q*^*D*^) or presence (*Q*^*Da*^) of acceptors, and the efficiency of energy transfer (or FRET efficiency, *E*) from donors to acceptors, namely the fractions of the number of excitations lost through radiative processes in the absence and presence of acceptors, and through energy transfer to the acceptor, respectively. In general, the quantum yield is defined as *the number of photons emitted divided by the total number of excitations*, while the FRET efficiency is defined as *the number of excitations transferred from donors to acceptors divided by the total number of excitations of the donor*. The total number of excitations equals the sum of excitations lost through different de-excitation pathways (regardless of whether the excitation light is continuous-wave or pulsed, or detection is with or without temporal resolution), and it may be computed by integrating Eqs. (1) over different time intervals and under different excitation conditions. Thus, expressions may be written for the donor and acceptor quantum yields in the absence of FRET,

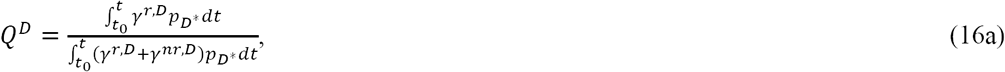

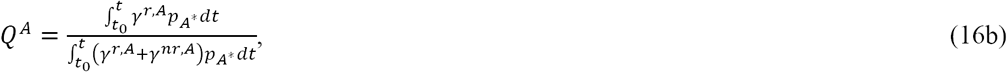

for the donor quantum yield in the presence of FRET,

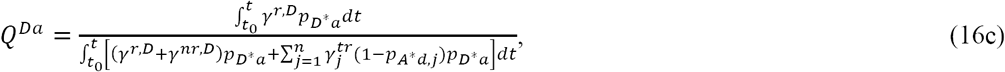

and the FRET efficiency,

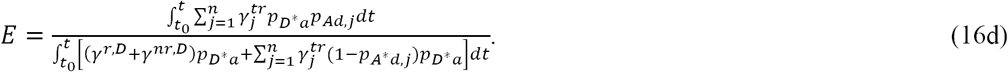

Since the rates of de-excitation are all constant, making use of the notations for the integrals of the probabilities from section II, these equations may be rewritten as

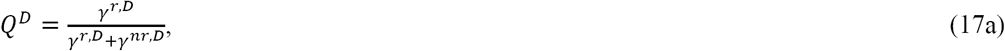

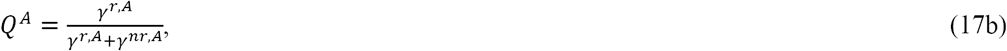

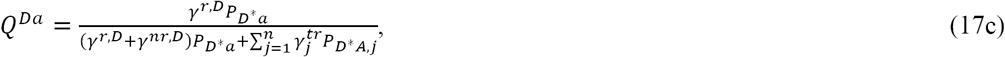

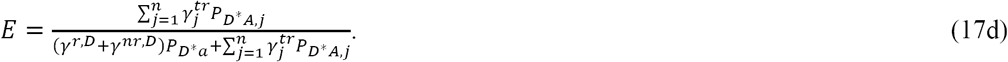

It is noteworthy that Eqs. (16c) and (16d) as well as (17c) and (17d) depend on the probabilities *P*_*D*A,j*_ and *P*_*D*a*_, which means that, in the most general case, they also depend to some extent on the excitation intensity as well as the extinction coefficients of the donors and acceptors. This is a departure from the widely used expressions for *Q*^*Da*^ and *E*, in which these probabilities are assumed to be equal to one another and therefore cancel out, leaving the two quantities dependent on de-excitation and transfer rates only. The net effect of such a dependence on experimental conditions is negligible, if the probability to find acceptors in their ground state, *p*_*Ad,j*_, is close to unity when the donor is in an excited state. This occurs when the acceptor is not directly excited by laser light. In that case, definitions (17c) and (17d) reduce themselves to the classical ones, according to which *Q*^*Da*^ and *E* depend only on the de-excitation rates (15). If, by contrast, the acceptor is also excited directly by laser light, as it is the case in many FRET approaches, the probability *p*_*Ad,j*_ is smaller than unity when the donor is in an excited state, which has some consequences, as detailed in sections II and III.

In either case, by rewriting Eq. (16d) as

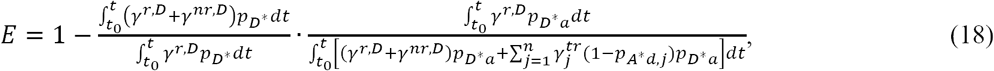

and using Eqs. (16a) and (16c) to replace the first and second terms, we obtain

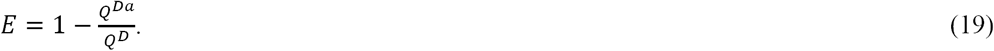

This is a fundamental relation of FRET (15), from which all expressions for the FRET efficiency, in terms of both temporally resolved and steady state fluorescence emission, may be derived, as it will be shown below.

#### II.4.2 Time-integrated fluorescence emission

Using Eqs. (4a) and (4b), the notations for the integrated probabilities used in Eqs. (4), and the definitions of quantum yields [Eqs. (17a,b)], we can now introduce the steady-state integrated emission of D or A per molecule in the absence of FRET:

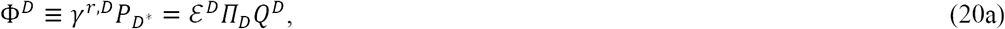

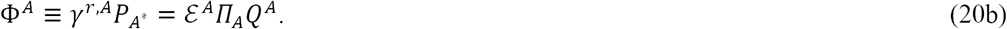

Successive use of Eqs. (4c) and (17c) gives the integrated emission per donor in the presence of FRET,

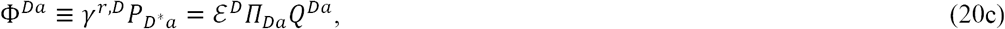

which, using Eq. (19), becomes

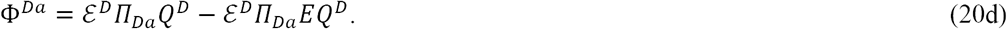

Finally, using successively Eqs. (4d), (17d), and (4c), we obtain for A in the presence of D,

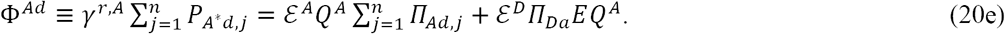

These equations will be used later on in this paper to derive various expressions for FRET efficiency as a function of measurable quantities.

#### II.4.3 Time-resolved fluorescence emission

In the absence of acceptor direct excitation, from Eqs. (11), we obtain for the emission intensities of the donors in the presence or absence of acceptors,

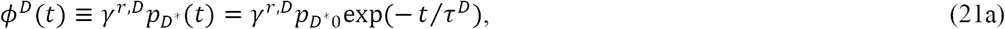

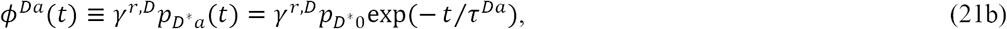

for temporal resolution higher than the fluorescence lifetime (i.e., *δt* ≫ τ). When Eqs. (21) are integrated over a time (*δt*) much longer than the lifetime of the excited state of the donor, but shorter than the repetition time of the laser pulses, we obtain

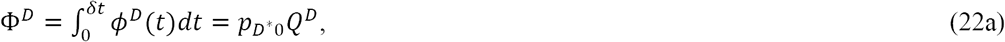

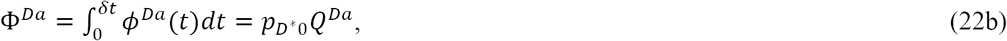

The results embodied by Eqs. (21) are customarily used to interpret the fluorescence decay of donors in FRET studies based on fluorescence lifetime imaging (FLIM). The results expressed by Eqs. (22) are formally similar to those of a steady-state situation [Eqs. (20)], whereby a continuous-wave light source is used and fluorescence emission is integrated over periods of time much longer than the excited lifetime of the donors and acceptors. This similarity provides a welcome consistency check of the kinetic theory outlined in this work.

Similarly, mathematical expressions may be obtained for the emission intensities of donors in the absence or presence of acceptors for the case of acceptors being directly excited by light,

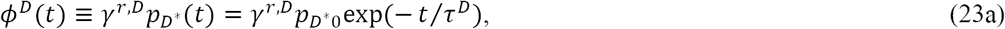

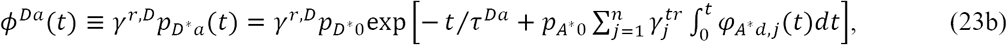

where τ^*Da*^ is as defined above – the donor lifetime in the presence of acceptors but in the absence of direct acceptor excitation. The integrals of these expressions depend on the particular form of φ_*A*d,j*_, as we have discussed in section II.

## III. RESULTS

### III.1. FRET efficiency expressions for pure oligomers

To obtain relationships between the efficiency of energy transfer and experimentally measured quantities, we rewrite Eq. (19) in different ways. By inserting the quantum yields of the donor in the presence and absence of FRET from Eqs. (20a) and (20c) into Eq. (19), we obtain,

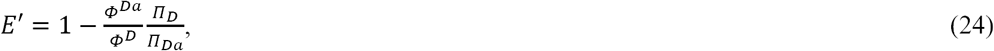

which for constant excitation intensity [see Eqs. (5)] becomes

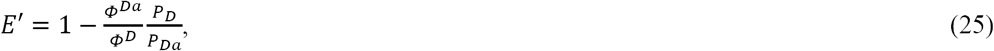

and for ultrashort-pulse excitation [see Eqs. (8)] becomes

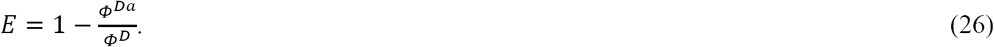

The last equation has been widely employed, together with various experimental schemes that allow the determination of Φ^*D*^ (e.g., acceptor photo-bleaching or use of two different excitation wavelengths (22, 24)), to compute the FRET efficiency from steady-state fluorescence intensities. Nevertheless, now we see that Eq. (26) should be used with caution, as it only applies to pulsed excitation and is expected to give somewhat erroneous results when used in conjunction with continuous-wave excitation, as is the case with confocal or wide-field microscopes. In addition, the use of photo-bleaching or two different excitation wavelengths to determine the donor emission in the absence of FRET leads to loss of pixel-level information in FRET imaging due to the fact that diffusion causes the molecular makeup of an image pixel to change during the long time it takes to acquire all the needed data.

To avoid the use multiple measurements or acceptor photo-bleaching, one can combine Eq. (20d) with the particular case of Eq. (20e) for the absence of acceptor direct excitation to obtain

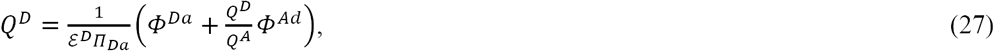

which inserted into Eq. (19) together with *Q*^*Da*^ from Eq. (20c), gives

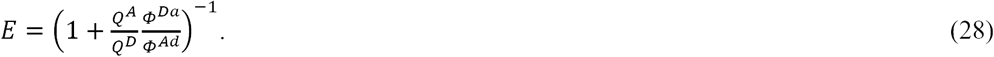

Equation (28) has been proposed previously and used successfully in many practical applications (10, 20, 28). Remarkably, it does not depend on probabilities and hence it is independent of the excitation level regardless of whether excitation is performed using CW or pulsed lasers. In addition, with this formula *E* may be determined upon a single excitation of the sample and does not rely on either multiple excitation wavelengths or acceptor photo-bleaching, as do other methods. This eliminates the need for strong approximations often used in FRET studies employing steady-state fluorescence emission and allows one to accurately compute *E*.

Using again the assumption that the acceptor is not directly excited by laser light (which leads to *p*_*Ad,j*_ = 1) and using the definitions of the lifetimes introduced by Eqs. (12), from Eq.(19) we obtain immediately:

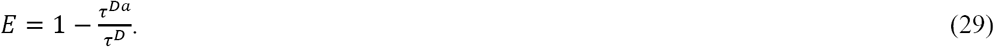

This equation is very popular with FLIM-FRET researchers, as it allows one to determine *E* from the fluorescence decay curves described theoretically by Eqs. (21) and which are obtained from FLIM experiments. Unfortunately, it is very difficult to extract with accuracy more than two different lifetimes from fluorescence decay curves, which makes this particular method applicable to probing interactions between only two molecules (i.e., dimers). This problem may be elegantly circumvented using the following approach.

When the acceptors are also directly excited by light, if the excitation light intensities are equal between the measurements of donors only and donors in the presence of acceptors, the denominators in Eqs. (16a) and (16c) cancel each other out once these expressions are inserted into Eq. (19). Thus, in our usual notations, we obtain:

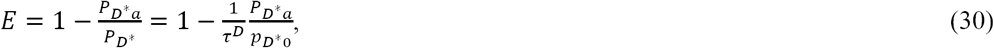

where *p*_*D\**0_ may be interpreted as the amplitude of the fluorescence decay curve (i.e., its height at *t* = 0). Further, by explicitly writing *P*_*D*a*_ as an integral, we obtain

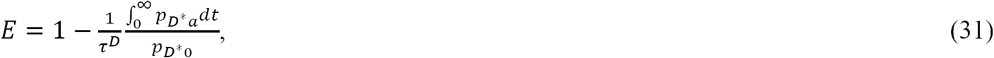

where *p*_*D*a*_ may be replaced by the fluorescence decay curve of the donor in the presence of acceptors, which may be then integrated numerically over the time interval between two excitation pulses to compute the FRET efficiency. By using the expression of *p*_*D*a*_ from Eq. (14), it may be easily seen that Eq. (31) reduces itself to Eq. (29) if there is no direct excitation of the acceptor (i.e., for *p*_*A*_*\**_0_= 0).

Equation (31) not only removes an approximation inherent in Eq. (3), but also promises to provide a means to compute the FRET efficiency for systems of oligomers of unknown size – and hence unknown number of lifetimes –, using time resolved fluorescence measurements. This feature could be exploited in the context of FRET spectrometry to determine the quaternary structure of macromolecules from distributions of FRET efficiencies obtained from pixel-level fluorescence measurements (20).

### III.2. FRET efficiency expressions for mixtures of oligomers and free monomers

#### III.2.1. Fluorescence of mixtures of interacting and non-interacting molecules

Up to this point in our derivations, fluorescence emission, whether expressed as photons emitted per unit of time (and denoted by *Ø*) or integrated over a longer time (denoted by upper case *Φ*), has implicitly referred to single molecules. This is indicated by the fact that we used probabilities of having the molecules in certain states, instead of concentrations of molecules in those states.

To describe emission intensities for ensembles of excited donors or acceptors, we multiply the previous results for *Φ* by the total concentrations of donors or acceptors in the sample volume, as appropriate. Additionally, we now consider the more general case that includes free donors and acceptors, as well as D-only and A-only oligomers, and we also replace the probabilities by concentrations. In light of these considerations and proceeding along the lines of reasoning surrounding Eqs. (19) from reference (15), Eqs. (20d) and (20e) are replaced by

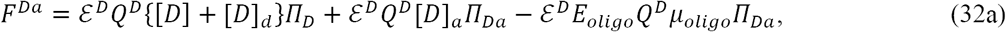

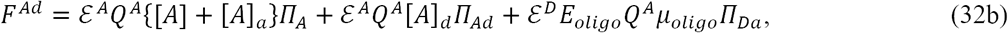

where [*D*]_*a*_ and [*A*]_*d*_ are the concentrations of donors in oligomers with acceptors, and the concentration of acceptors within oligomers with donors, respectively, the term ε^*D*^*Q*^*D*^{[*D*] + [*D*]_*d*_}П_*D*_ accounts for the emission of free donors and donors in complexes with other donors (but no acceptors), and ε^*A*^*Q*^*A*^{[*A*] + [*A*]_*a*_} is the emission of free acceptors and acceptors in complexes with other acceptors (but no donors).

To simplify these expressions, we introduce the notations

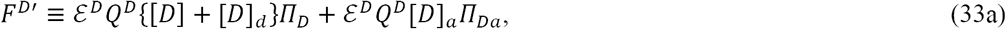

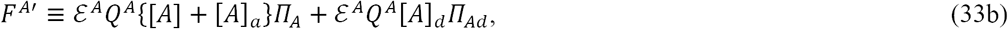

for the fluorescence of donors and acceptors, and

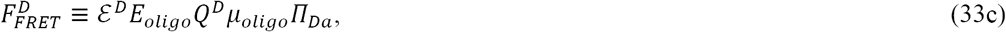

and

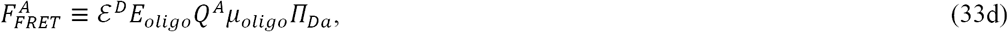

for the loss or gain due to FRET. We also introduce the concept of *apparent FRET efficiency, E*_*app*_, of a mixture of free as well as associated donors, some of which may be involved in FRET with acceptors, which is expressed by (12, 15):

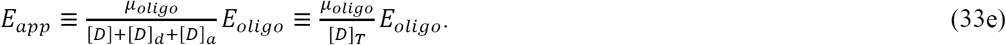

This is the quantity of interest in FRET experiments and provides the connection to the oligomeric size and configuration (i.e., quaternary structure) (20).

With these notations and definitions, Eqs. (32) become

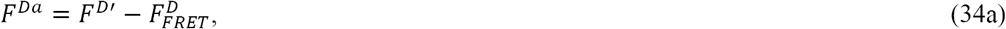

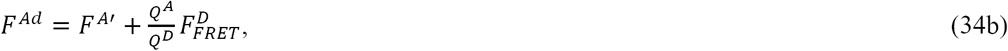

as it has already been proposed before (10, 15). Note that we used “prime” to denote *F*^*D*′^ and *F*^*A*′^, because, according to Eqs. (33a) and (33b), they are only equal to the donor-only and acceptor-only fluorescence emissions if the probabilities follow the relations: П_*D*_ = П_*Da*_, and П_*A*_ = П_*Ad*_. In that case, Eqs. (33a) and (33b) become

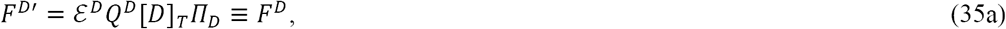

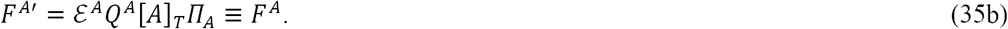

The generally accepted definition of FRET efficiency in terms of integrated emission intensities (over entire emission spectra), by analogy to Eq. (26), is given by the equation:

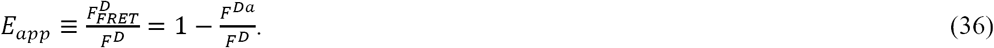

If the donor and acceptor are chosen such that the acceptor is not directly excited by light at a wavelength at which the donor is excited, then *F*^*A*′^ = 0, and combination of Eqs. (34) gives

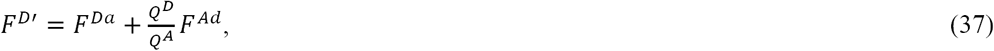

which is equal to *F*^*D*^ for pulsed excitation light, but not for CW excitation. Inserting Eq. (37) into Eq. (36), we obtain an equation,

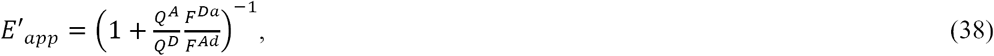

which has a similar form to Eq. (28), which applies to pure systems of oligomers.

Since we used *F*^*D*′^ and not *F*^*D*^ to derive Eq. (38), it is necessary to verify that it actually gives the correct result. By inserting Eqs. (34) (with *F*^*A*′^ = 0) into the right-hand-side of Eq. (38), this equation reduces itself to the correct expression, 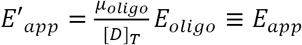, but only for pulsed and not for CW excitation, since in the former case П_*Da*_ = П_*D*_ = 1, as mentioned above and further discussed below.

When *γ*^*ex,A*^ ≠ 0, we have *F*^*A*^ ≠ 0, and Eqs. (34) may be solved only if a second excitation wavelength is used, since the system of Eqs. (34) is otherwise underdetermined. This problem has been previously tackled and its results successfully applied to probing oligomerization of membrane receptors (11, 12, 14).

#### III.2.2 Determination of E_app_ and concentrations using two excitation wavelengths

If, in addition to the donor concentration and the FRET efficiency, the acceptor concentration also needs to be determined (11, 14), the theory of the method has to incorporate the direct excitation of the acceptor as well as additional equations by adding a second excitation wavelength.

In the case of two excitation wavelengths, one may write the following variants of Eqs.(34) for the donor and acceptor emission in the presence of FRET:

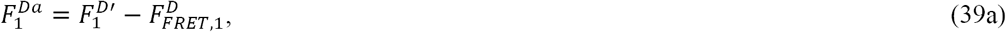

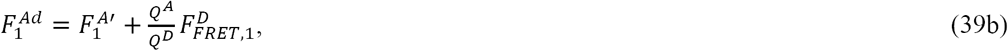

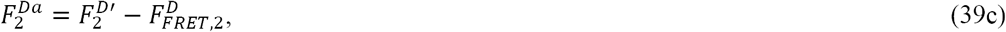

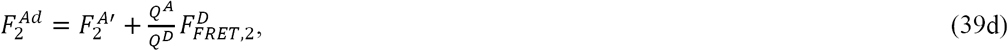

where the subscripts “1” and “2” stand for the first and second excitation wavelength, respectively. In addition, using the notations given by Eqs. (33), we introduce the following notations for the ratios of the various terms in Eqs. (39):

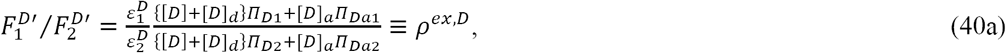

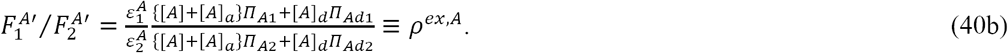

For mixtures of molecules in different oligomeric states, ρ^*ex,D*^ and ρ^*ex,A*^ may be easily determined, though only for the case of pulsed excitation (for which П_*D*_ = П_*Da*_ = П_*A*_ = П_*Ad*_ =1, and therefore 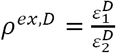 and 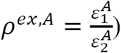, by measuring separately the relative fluorescence emission following excitation at the two wavelengths of samples containing only donors or only acceptors.

After dividing Eq. (39a) by *Q*^*D*^ and (39b) by *Q*^*A*^ and adding up the resulting expressions, we have:

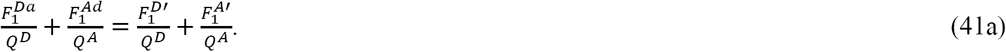

Similarly, we obtain the following expression by combining Eqs. (39c) and (39d):

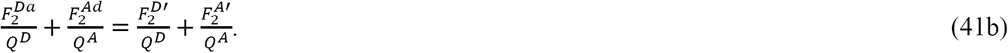

Then, substituting 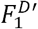 and 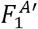 from Eqs. (40a) and (40b), respectively, into Eq. (41a), and dividing the resulting equation by ρ^*ex,D*^ we obtain

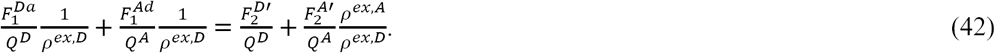

Subtracting Eq. (42) from Eq. (41b) and rearranging the terms, we obtain:

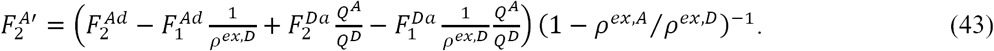

For pulsed excitation, П_*D*_ = П_*Da*_ = 1 and therefore Eqs. (32a) and (40a) provide that 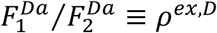, in which case Eq. (43) becomes

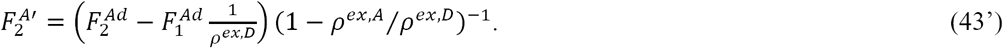

Further, by solving Eq. (41a) for 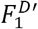 and using Eq. (40b) to substitute for 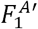, we obtain

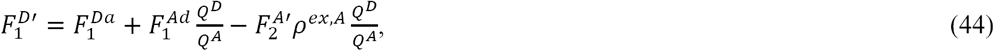

where 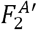 is determined from experiments via Eq. (43).

Finally, by inserting 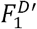 from Eq. (44) into Eq. (36), we obtain:

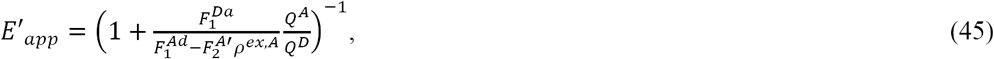

where 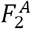 is connected to experiment via Eq. (43). For pulsed excitation, we may substitute 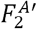 from Eq. (43’) and obtain:

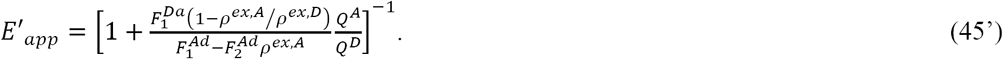

Equation (45’) as well as similar or approximate forms of it have been used previously by us or other researchers (11, 12, 17, 24) to determine the FRET efficiency for pure or mixed forms of molecular complexes of different sizes (11, 24). By inserting the particular forms taken by Eqs. (32) and (33b) for П_*D*_ = П_*Da*_ = П_*A*_ = П_*Ad*_ = 1 into the right-hand-side of Eq. (45’), we find that this equation reduces itself to the correct expression, 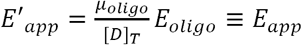. By contrast, the more general Eq. (45) combines integrated probabilities of donors and acceptors to be in excited states for two different wavelengths – all of which take unknown values. Therefore, for CW excitation, the FRET efficiency is subject to systematic errors. These errors could be corrected for (see the Supplementary materials), but that process requires time-resolved measurements in addition to CW-based measurements.

## IV. DISCUSSION

### IV.1. Determination of *E* from time-resolved measurements

As discussed above, if the acceptor is already in an excited state when the donor is excited, as a result of acceptor direct excitation by laser light, quantum mechanics rules prevent the donor from transferring its excitation to the acceptor. The magnitude of this effect is estimated by evaluating the integral in the exponent of Eq. (14) using the approximations that the acceptor fluorescence follows the same exponential decay curve as it would in the absence of FRET, φ_*A*d,j*_*(t)* = exp(−*t/*τ^*A*^), and that all acceptors are equally excited, whether by laser light or via FRET. As shown in Figure S1 of the Supplementary Results, the change in the donor excited lifetime is rather small, even for tetramers consisting of three acceptors and one donor. To estimate the contribution of the acceptor direct excitation to *E*, assuming again that the acceptor fluorescence follows an exponential decay, after performing the integration in the exponent of Eq. (14) and inserting the resulting probability into Eq. (31), we obtain:

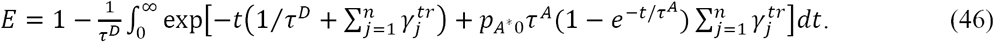

By inserting into this expression the parameter values used for simulating the fluorescence decay curves in Figure S2, we obtain *E* = 0.783 in the absence and *E =* 0.768 in the presence of 10% acceptor direct excitation, respectively. This overestimate of *E* when acceptor direct excitation is taken into account is rather modest (∼2%), although it may play some role in the interpretation of the results from high-precision experiments.

Regardless of whether the competition between energy transfer and laser light for exciting the acceptors produces measurable effects, Eq. (31) suggests a simple alternative way of computing the FRET efficiency in mixtures of monomers and oligomers with different proportions of donors and acceptors, which are expected to generate a superposition of several exponential decay curves with different lifetimes. As we have mentioned above, such situations may not be tackled using Eq. (29). Even if it were possible to extract several lifetimes from a fluorescence decay curve, one would need to know a priori the number and sizes of the different types of oligomers present (in order to know how many lifetimes to extract), which is the very piece of information that one actually needs to extract from FLIM experiments. This creates a vicious circle, which may be avoided by integrating the fluorescence decay curves – rather than resolving them into multiple exponentials –, and then using Eq. (31) to compute the FRET efficiency.

This proposed method, which is different from FLIM and which we tentatively name “time-resolved intensity measurements” (TRIM), should be useable in conjunction with FRET spectrometry, which allows determination of quaternary structures but has so far been implemented using average-intensity-based fluorescence measurements only (20, 28).

### IV.2. Estimating and testing systematic errors introduced by CW excitation

We have suggested in section IV that assuming that the probability of the donors to be in their ground state in the absence of FRET is equal to that corresponding to the presence of FRET – i.e., П_*D*_ = П_*Da*_ (or *P*_*D*_ = *P*_*Da*_) –, may lead to errors in computing the FRET efficiency using the standard expressions for FRET efficiency and CW excitation light sources. To estimate the errors in the FRET efficiency when using Eq. (25) for systems of pure oligomeric complexes, we first rewrite Eq. (25) in terms of the true FRET efficiency, *E*, given by Eq. (26), as

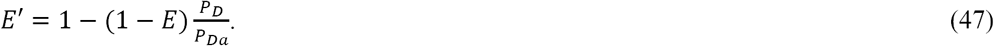

Replacing the integrated probabilities by their corresponding expressions [Eqs. (S1) and (S2)] derived in Supplementary Results, Eq. (47) becomes:

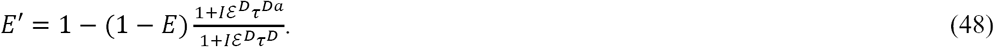

Assuming that a laser beam with an average power of 1 mW and a wavelength of 433 nm is focused to a diffraction limited spot of radius 200 nm onto a sample containing dimers consisting of single Cerulean (30) fluorescent proteins (with extinction coefficient ε^*D*^ = 4,300 M^−1^ m^−1^) fused to some other fluorescent protein to form FRET dimers, the excitation rate for a donor molecule given by Eq. (2) is *γ*^*ex,D*^ = I*ε*^*D*^ = 2.85 × 10^8^ s^−1^. The lifetime of Cerulean in the absence of FRET is approximately 3.2 ns (31). Its lifetime in the presence of FRET with a true FRET efficiency value of, let us say, *E* = 0.28 is estimated from Eq. (29) to be 2.3 ns. By inserting these values into Eq. (48), we obtain a FRET efficiency *E*′ = 0.38, which is some 34 % larger than the true FRET efficiency, *E,* of our assumed dimers. The excitation power of 1 mW used in our estimates above is not atypical in experiments involving fluorescent molecules detection (32), and so this kind of errors in the FRET efficiency could not be considered as exceptional. Nevertheless, even if the excitation power decreased by an order of magnitude to 0.1 mW, the error in FRET efficiency predicted by Eq. (48) for CW excitation would still be 6%.

Systematic errors in FRET efficiency measurements based on CW excitation light have been noticed ever since Stryer’s introduction of the “spectrometric ruler” (33). In a somewhat more recent paper, Deniz and co-workers (34) used di-nucleotides of the same lengths labelled fluorescently at different positions, such that the tags were positioned at various distances from one another, and performed single particle FRET measurements using CW excitation. By plotting the FRET efficiency values versus distance, Deniz et al noticed a shift towards larger distances in the experimental data plot compared to the plot predicted by Förster’s well-known formula. In light of our discussion above, it is reasonable to assume that least part of those discrepancies arose from implicit approximations concerning the integrated probabilities. To test that hypothesis, we substitute τ^*Da*^ from Eq. (29) into Eq. (48) to obtain a relationship,

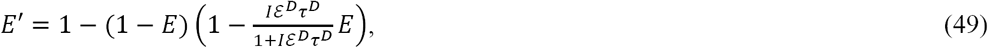

that connects the measured efficiency *E’* to that predicted by Förster’s formula for different values of the inter-fluorophore distance, *R*. By reanalyzing Deniz et al’s data, we found that the *E’* vs. *R* curve predicted by Eq. (49) is significantly shifted to the right and better fits the experimental data for the same Förster distance (*R*_*0*_ = 65 Å) as in the work described above (see Supplementary Figure S2). This confirms the validity of the theory described in this work.

As for the systematic errors affecting the output of Eq. (38) when using CW excitation, we could only infer that they will be at least as large as those estimated in the paragraph above. For reference, we estimate the values of the four integrated probabilities, using the expressions provided in the Supplementary Results section, for the experimental conditions described above and, in addition, assuming that the acceptors consist of Venus fluorescent molecules (35) with an extinction coefficient ε^*A*^ = 9,200 M^−1^ m^−1^ and a lifetime of 3 ns. For 1 ms integration time and 1 mW of excitation power, these values are, *P*_*D*_= 5.2 × 10^−4^ s, *P*_*Da*_= 6.0 × 10^−4^ s, *P*_*A*_= 3.5 × 10^−4^ s, *P*_*Ad*_= 2.2 × 10^−4^ s, with the donor integrated probability being 13% smaller and the acceptor integrated probability 58% larger in the presence of FRET compared to their respective values in the absence of FRET. As seen, the acceptor integrated probability is lower in the presence of FRET, compared to that in the absence of FRET, because excitation through both FRET and laser light causes the acceptors to spend less time in their ground state. Since, in the expressions for *F*^*Da*^ and *F*^*Ad*^ in Eqs. (32), each such integrated probability multiplies the concentration of a different kind of donor or acceptor (i.e., free, as well as bound to their own or different kind of molecule), taking the ratio of *F*^*Da*^ and *F*^*Ad*^ to compute the FRET efficiency using Eq. (38) would lead to different values of *E’*_*app*_ depending on the relative proportion of the differing fluorescent species, with none of these values being actually equal to *E’*_*app*_.

Following the same kind of reasoning as above, as well as the arguments made in section IV, it may be concluded that for CW excitation the use of Eq. (45) also leads to systematic errors in the FRET efficiency for mixtures of molecules in different states of association. Furthermore, the same kind of errors affect the determination of the concentrations of donors and acceptors using Eqs. (43) and (44).

### IV.3. Avoiding systematic and random errors in *E*_*app*_ when using pulsed excitation

As mentioned above, for pulsed excitation light, Eqs. (38) and (45’) correctly lead to 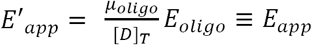 and are expected to provide values of FRET efficiency with arbitrary accuracy, sine П_*Da*_ = П_*D*_ = 1, as long as its underlying assumption of no acceptor direct excitation is valid. Therefore, proper use of Eqs. (38) and (45’) allows one to determine the quaternary structure of proteins using FRET spectrometry, which relies on plotting histograms of frequencies of *E*_*oligo*_ values for systems of oligomers with different combinations of donors and acceptors (20). Additionally, it is possible to determine the donor concentration from Eq. (37), using the same measurement performed for determining *E*_*app*_, as proposed previously (36).

Nevertheless, either one of those equations may be affected by systematic errors, depending on the particular experimental protocol used. For instance, Eq. (38) could overestimate the FRET efficiency if the acceptor is directly excited by light to a significant degree. At the same time, acceptor or donor photo-bleaching or photo-switching (37) could cause 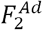, ρ^*ex,A*^, and ρ^*ex,D*^ to change relative to the absence of photochemical effects. Specifically, if the excitation at the first wavelength leads to photo-bleaching of a fraction of donors or acceptors, the apparent concentration of the molecules detected upon excitation at the second wavelength scan decreases. To take that into account, for pulsed excitation (i.e., П_*D*_ = П_*Da*_ =П_*A*_ = П_*Ad*_ = 1), using Eq. (32b), we rewrite 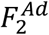 in Eq. (45’) as

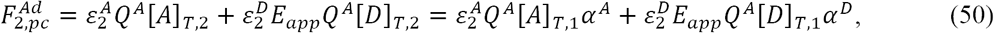

where the subscript “pc” stands for “photochemical” effects (which may include photo-bleaching and photo-switching), *α*^*A*^ = [*A*]_*T*,2_*/*[*A*]_*T*,1_, and *α*^*D*^ = [*D*]_*T*,2_*/*[*D*]_*T*,1_.

When the determination of the ρ^*ex,A*^ and ρ^*ex,D*^ values is done using the same sample (e.g., cells), these quantities too will be affected by photo-bleaching (or photo-switching), as illustrated by the following forms of Eqs. (40) for the case of pulsed excitation:

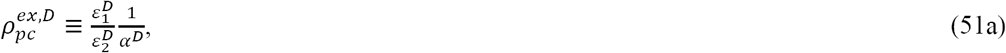

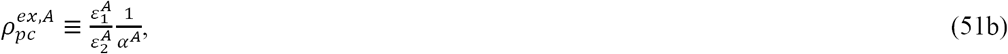

which depend on *α*^*A*^ and *α*^*D*^. By inserting Eqs. (5) and (51) into Eq. (45’) and also using other relationships provided above as needed, it may be easily seen that *α*^*A*^ and *α*^*D*^ cancel out in Eq. (45’) and this equation still reduces itself to *E*′_*app*_ = *E*_*app*_, as it is desirable.s

By contrast, when the FRET measurements are performed on cells, where the molecules are less free to move than in a purely aqueous solution and hence are more susceptible to photo-bleaching or photo-switching, while the determination of ρ^*ex,A*^ and ρ^*ex,D*^ is based on measurements of pure solutions, in which molecules diffuse freely (and therefore are less prone to photo-bleaching or photo-switching), Eq. (45’) will depend on *α*^*A*^ and *α*^*D*^, and hence the final computed FRET efficiency will be affected by the photochemical effects. This may explain why the validity of the kinetic theory of FRET could not be confirmed in previous work (38).

It is also possible that the effect of photo-bleaching is seen even during the first excitation scan, which could in principle affect the output of both Eq. (38) and (45’). For pulsed excitation (i.e., П_*D*_ = П_*Da*_ = П_*A*_ = П_*Ad*_ = 1), we have

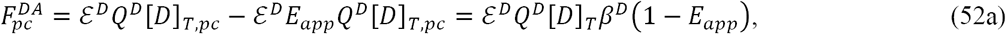

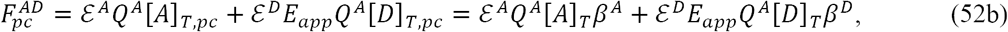

where *β*^*A*^ = [*A*]_*T,pc*_*/*[*A*]_*T*_, and *β*^*D*^ = [*D*]_*T,pc*_*/*[*D*]_*T*_. In the absence of direct acceptor excitation (i.e., for *ε*^*A*^ = 0), insertion of Eqs. (52) into Eq. (38) leads correctly to *E*′_*app*_ = *E*_*app*_, because *β*^*D*^ appears both in the denominator and the numerator and therefore cancels out. However, if Eqs. (52) are inserted into Eq. (45’), for which *ε*^*A*^ may be different from zero, the beta correction factors cancel out only if the calibration of ρ^*ex,A*^ and ρ^*ex,D*^ is performed using the same kind of sample, as in the case of the alpha correction factors. Therefore, Eq. (45’) may be affected by additional systematic errors, depending on the particular type of calibration used, when compared to Eq. (38).

The only systematic error affecting Eq. (38) that we could conceive of is the possibility for the acceptors to be directly excited by light to a small extent. This is probably why in a previous publication (29), any discrepancy between experiment and the kinetic theory of FRET was seen to be small (∼ 4%, see appendix C in that reference) compared to the discrepancy of ∼ 15% registered in earlier work when an equation similar to Eq. (45’) was used. Errors possibly affecting both approaches may exist, such as random errors and errors caused by sample inhomogeneity (19).

### IV.4. Optimization of the protocols for determination of molecular concentrations

Our analysis above has already revealed optimal strategies for determination of the FRET efficiency using appropriate excitation wavelengths, equations, and calibration strategies, that are both simple enough and free from systematic errors. At this point, it remains to identify robust methods for determining the total concentrations of molecules.

The first method we identified involves determining the donor concentration from a first scan of the sample using an excitation wavelength that does not produce any significant excitation of the acceptor, and determining the acceptor concentration from a second scan at an excitation wavelength that excites the acceptor most efficiently. A theoretical expression for the donor concentration in the case of pulsed excitation light in the absence of acceptor direct excitation is obtained, by combining Eqs. (35a) and (37), as

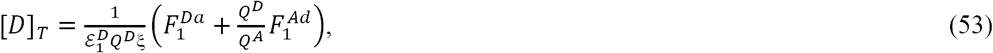

in which the theoretical brightness (i.e., *ε*^*D*^*Q*^*D*^) has been multiplied by an instrumental factor, ξ, which accounts for detection laser excitation power, detection sensitivity, etc. An expression for the acceptor concentration is given by the equation

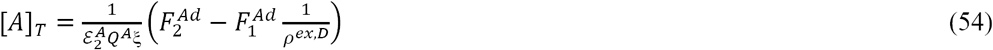

which was obtained by dividing Eq. (43’) by the effective acceptor brightness (including the instrumental factor ξ) and assuming that ρ^*ex,A*^ = 0. The total concentration in this case is obtained as the sum between the donor and acceptor concentrations.

The second method for determining the total concentration of donor and acceptor molecules has been suggested recently in a different context (27). In this method, the second excitation wavelength is chosen such that the excitation rates of the donor and acceptor are equal (i.e., 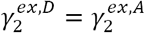, which is equivalent to 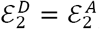). With this, we obtain the following expression by dividing Eq. (39c) by 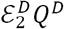 and Eq. (39d) by 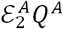 and adding the resulting expressions side by side:

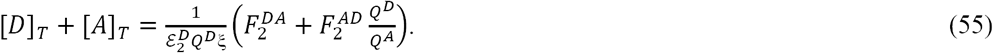

For all equations in this paragraph, the effective brightness (*ε*^*D*^*Q*^*D*^ξ or *ε*^*A*^*Q*^*A*^ξ) may be determined experimentally using either calibration against fluorescent molecule solutions with known concentrations (12) or, more elegantly and precisely, using fluorescence fluctuation analysis (39-41) of cells expressing monomeric constructs (42).

## V. CONCLUSION

We have derived equations expressing the fluorescence emission of acceptors and donors in the presence of FRET [i.e., Eqs. (33)] rigorously from the kinetic model of FRET using certain assumptions regarding the de-excitation rates and probabilities of fluorescent molecules to be in their ground or excited sates. We have found that those assumptions are not obeyed automatically and require deliberate decisions by the experimentalist in using one expression or the other, which is equivalent to choosing different sample excitation and detection conditions. Under those same assumptions, some known as well as some previously unknown expressions are rigorously derived, which link the FRET efficiency to experimentally measurable parameters, such as fluorescence lifetimes or integrated fluorescence intensities. Also interestingly, it was found that in situations where excitation is produced by continuous wave sources, significant systematic errors are introduced in the computation of FRET efficiency as well as concentrations of molecules. By contrast, pulsed light sources, whether or not used in conjunction with temporal resolution for detection of the fluorescence, provide means of extracting the values of all of these quantities free from significant systematic errors. Nevertheless, systematic errors are still possible if calibrations are not performed carefully or inadequate equations are used for given experimental conditions.

Based on the present analysis, we are able to suggest an optimal strategy for quantitative FRET spectrometric investigations. In the case of no temporal resolution, the following procedure is recommended.

(i) Use pulsed excitation to avoid systematic errors caused by the dependence of all equations on integrated probabilities (and, hence, on concentrations).
(ii) Use spectral resolution and unmixing to separate donor and acceptor signals from each sample scan.
(iii) Perform a “*FRET scan”* of the sample at an excitation wavelength at which the acceptor is minimally excited by light while the donor is excited maximally and use Eq. (38) to compute the FRET efficiency as well as the donor concentration [Eq. (44) with ρ^*ex,A*^ = 0]. If there is any direct excitation of acceptors by light and low errors are essential, scan the sample at a second wavelength and use Eq. (45’) to perform careful corrections as described in the discussion section.
(iv) (***a***) Perform a “*concentration scan*” of the sample at an excitation wavelength at which mostly the acceptor is excited and use Eq. (43’), assuming again ρ^*ex,A*^ = 0, to compute the concentration of acceptors. The total concentration is computed as the sum of [*D*]_*T*_ and [*A*]_*T*_. (***b***) Alternatively, perform a *concentration scan* at a wavelength at which the donors and acceptors are excited equally well (i.e., 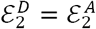), and use Eq. (55) to compute the total concentration.

Some of the precautions outlined above have already been used to determine quaternary structures of proteins as well as the relative proportion of the differing structures using large numbers of acquired cellular images. When combined with certain experimental tools, this approach will open the way for combining FRET spectrometry with FRET stoichiometry measurements for each image pixel in the future (20, 28).

As suggested in the previous sections, it should also be possible to devise a FRET spectrometry method based on time resolved measurements, in which analysis should be done by integrating rather than unmixing the fluorescence decay curves into different lifetimes. In that case, Eq. (31) would have to be used for calculation of the FRET efficiency, while molecular concentrations could still rely on the same equations used in the case of no temporal resolution. This idea remains to be tested experimentally in the future.

## Acknowledgments

The author thanks Dr. Steven Vogel (NIH) for several useful discussions regarding tests of the kinetic theory, Drs. Gabriel Biener and Michael Stoneman for useful discussions regarding some of the derivations presented in this article, and to Dhruba Adhikari, Dammar Badu, Gabriel Biener, and Joel Paprocki for critical reading of the manuscript. This work was partly funded by a National Science Foundation grant (PHY-1626450) and the UWM Research Growth Initiative grants 101X333 and 101X340.

## Footnotes

1 The sifting property of the Dirac delta function provides that 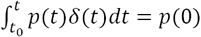 for *t*_0_< 0 < *t*, and 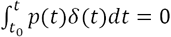 for any other integration limits.

## REFERENCES

1. Lakowicz, J. R. 2006. Principles of Fluorescence Spectroscopy. Springer, New York.

2. Clegg, R. M. 1996. Fluorescence Resonance Energy Transfer. In Fluorescence Imaging Spectroscopy and Microscopy. X. F. Wang, and B. Herman, editors. Wiley-Interscience, New York.

3. Selvin, P. R. 2000. The renaissance of fluorescence resonance energy transfer. Nat Struct Biol 7:730–734.

4. Meyer, B. H., J.-M. Segura, K. L. Martinez, R. Hovius, N. George, K. Johnsson, and H. Vogel. 2006. FRET imaging reveals that functional neurokinin-1 receptors are monomeric and reside in membrane microdomains of live cells. Proc. Natl. Acad. Sci. U.S.A. 103:2138–2143.

5. Onuki, R., A. Nagasaki, H. Kawasaki, T. Baba, T. Q. Uyeda, and K. Taira. 2002. Confirmation by FRET in individual living cells of the absence of significant amyloid beta-mediated caspase 8 activation. Proc Natl Acad Sci U S A 99:14716–14721.

6. Li, Y. C., R. V. Shivnaraine, D. D. Fernandes, H. Q. Ji, F. Huang, J. W. Wells, and C. C. Gradinaru. 2014. Nature of the M-2 Muscarinic Receptor Signaling Complex Revealed by Dual-Color FCS and FRET. Biophysical Journal 106:101a–101a.

7. Flynn, D. C., A. R. Bhagwat, M. H. Brenner, M. F. Nunez, B. E. Mork, D. W. Cai, J. A. Swanson, and J. P. Ogilvie. 2015. Pulse-shaping based two-photon FRET stoichiometry. Optics Express 23:3353–3372.

8. Singh, D. R., F. Ahmed, S. Sarabipour, and K. Hristova. 2017. Intracellular Domain Contacts Contribute to Ecadherin Constitutive Dimerization in the Plasma Membrane. Journal of Molecular Biology 429:2231–2245.

9. Li, M., L. G. Reddy, R. Bennett, N. D. Silva, L. R. Jones, and D. D. Thomas. 1999. A fluorescence energy transfer method for analyzing protein oligomeric structure: application to phospholamban. Biophys. J. 76:2587–2599.

10. Raicu, V., M. R. Stoneman, R. Fung, M. Melnichuk, D. B. Jansma, L. F. Pisterzi, S. Rath, M. Fox, J. W. Wells, and D. K. Saldin. 2009. Determination of supramolecular structure and spatial distribution of protein complexes in living cells. Nature Photonics 3:107–113.

11. King, C., M. Stoneman, V. Raicu, and K. Hristova. 2016. Fully quantified spectral imaging reveals in vivo membrane protein interactions. Integrative Biology 8:216–229.

12. Stoneman, M. R., J. D. Paprocki, G. Biener, K. Yokoi, A. Shevade, S. Kuchin, and V. Raicu. 2017. Quaternary structure of the yeast pheromone receptor Ste2 in living cells. Biochimica Et Biophysica Acta-Biomembranes 1859:1456–1464.

13. Corby, M. J., M. R. Stoneman, G. Biener, J. D. Paprocki, R. Kolli, V. Raicu, and D. N. Frick. 2017. Quantitative microspectroscopic imaging reveals viral and cellular RNA helicase interactions in live cells. Journal of Biological Chemistry 292:11165–11177.

14. Mishra, A. K., M. Gragg, M. Stoneman, G. Biener, J. A. Oliver, P. Miszta, S. Filipek, V. Raicu, and P. Park. 2016. Quaternary structures of opsin in live cells revealed by FRET spectrometry. Biochemical Journal.

15. Raicu, V. 2007. Efficiency of Resonance Energy Transfer in Homo-Oligomeric Complexes of Proteins. Journal of Biological Physics 33:109–127.

16. King, C., V. Raicu, and K. Hristova. 2017. Understanding the FRET Signatures of Interacting Membrane Proteins. Journal of Biological Chemistry 292:5291–5310.

17. Thaler, C., S. V. Koushik, P. S. Blank, and S. S. Vogel. 2005. Quantitative Multiphoton Spectral Imaging and Its Use for Measuring Resonance Energy Transfer. Biophys J 89:2736–2749.

18. Biener, G., M. R. Stoneman, G. Acbas, J. D. Holz, M. Orlova, L. Komarova, S. Kuchin, and V. Raicu. 2014. Development and Experimental Testing of an Optical Micro-Spectroscopic Technique Incorporating True Line-Scan Excitation. Int J Mol Sci 15:261– 276.

19. Singh, D. R., and V. Raicu. 2010. Comparison between whole distribution- and average-based approaches to the determination of fluorescence resonance energy transfer efficiency in ensembles of proteins in living cells. Biophys J 98:2127–2135.

20. Raicu, V., and D. R. Singh. 2013. FRET Spectrometry: A New Tool for the Determination of Protein Quaternary Structure in Living Cells. Biophysical Journal 105:1937–1945.

21. Elangovan, M., R. N. Day, and A. Periasamy. 2002. Nanosecond fluorescence resonance energy transfer-fluorescence lifetime imaging microscopy to localize the protein interactions in a single living cell. J Microsc 205:3–14.

22. Jares-Erijman, E. A., and T. M. Jovin. 2003. FRET imaging. Nat Biotech 21:1387–1395.

23. Neher, R. A., and E. Neher. 2004. Applying spectral fingerprinting to the analysis of FRET images. Microsc. Res. Tech. 64:185–195.

24. Raicu, V., D. B. Jansma, R. J. Miller, and J. D. Friesen. 2005. Protein interaction quantified in vivo by spectrally resolved fluorescence resonance energy transfer. Biochem J 385:265–277.

25. Vogel, S. S., C. Thaler, and S. V. Koushik. 2006. Fanciful FRET. Sci STKE 2006:re2.

26. Singh, D. R., M. M. Mohammad, S. Patowary, M. R. Stoneman, J. A. Oliver, L. Movileanu, and V. Raicu. 2013. Determination of the quaternary structure of a bacterial ATP-binding cassette (ABC) transporter in living cells. Integrative Biology 5:312–323.

27. Raicu, V. 2018. Extraction of information on macromolecular interactions from fluorescence micro-spectroscopy measurements in the presence and absence of FRET. Spectrochimica Acta Part A: Molecular and Biomolecular Spectroscopy 199:8.

28. Raicu, V., and W. F. Schmidt. 2017. Advanced Microscopy Techniques. In G-Protein-Coupled Receptor Dimers. K. Herrick-Davis, G. Milligan, and G. Di Giovanni, editors. Humana Press. 39–75.

29. Patowary, S., L. F. Pisterzi, G. Biener, J. D. Holz, J. A. Oliver, J. W. Wells, and V. Raicu. 2015. Experimental verification of the kinetic theory of FRET using optical microspectroscopy and obligate oligomers. Biophysical Journal 108:1613–1622.

30. Rizzo, M. A., G. H. Springer, B. Granada, and D. W. Piston. 2004. An improved cyan fluorescent protein variant useful for FRET. Nature Biotechnology 22:445–449.

31. Sarkar, P., S. V. Koushik, S. S. Vogel, I. Gryczynski, and Z. Gryczynski. 2009. Photophysical properties of Cerulean and Venus fluorescent proteins. J Biomed Opt 14:034047.

32. Nie, S. M., D. T. Chiu, and R. N. Zare. 1994. Probing Individual Molecules with Confocal Fluorescence Microscopy. Science 266:1018–1021.

33. Stryer, L., and R. P. Haugland. 1967. Energy transfer: a spectroscopic ruler. Proc Natl Acad Sci U S A 58:7.

34. Deniz, A. A., M. Dahan, J. R. Grunwell, T. J. Ha, A. E. Faulhaber, D. S. Chemla, S. Weiss, and P. G. Schultz. 1999. Single-pair fluorescence resonance energy transfer on freely diffusing molecules: Observation of Forster distance dependence and subpopulations. Proceedings of the National Academy of Sciences of the United States of America 96:3670–3675.

35. Nagai, T., K. Ibata, E. S. Park, M. Kubota, K. Mikoshiba, and A. Miyawaki. 2002. A variant of yellow fluorescent protein with fast and efficient maturation for cell-biological applications. Nat Biotechnol 20:87–90.

36. Mishra, A. K., T. Mavlyutov, D. R. Singh, G. Biener, J. Yang, J. A. Oliver, A. Ruoho, and V. Raicu. 2015. The sigma-1 receptors are present in monomeric and oligomeric forms in living cells in the presence and absence of ligands. Biochemical Journal 466:263–271.

37. Markwardt, M. L., G. J. Kremers, C. A. Kraft, K. Ray, P. J. C. Cranfill, K. A. Wilson, R. N. Day, R. M. Wachter, M. W. Davidson, and M. A. Rizzo. 2011. An Improved Cerulean Fluorescent Protein with Enhanced Brightness and Reduced Reversible Photoswitching. Plos One 6.

38. Koushik, S. V., P. S. Blank, and S. S. Vogel. 2009. Anomalous surplus energy transfer observed with multiple FRET acceptors. PLoS ONE 4:e8031.

39. Qian, H., and E. L. Elson. 1990. On the Analysis of High-Order Moments of Fluorescence Fluctuations. Biophysical Journal 57:375–380.

40. Chen, Y., J. D. Muller, S. A. Sanchez, G. Patterson, D. W. Piston, and E. Gratton. 2000. Towards quantitative measurement of GFP in vivo with fluorescence fluctuation spectroscopy. Biophysical Journal 78:280a–280a.

41. Godin, A. G., S. Costantino, L.-E. Lorenzo, J. L. Swift, M. Sergeev, A. Ribeiro-da-Silva, Y. De Koninck, and P. W. Wiseman. 2011. Revealing protein oligomerization and densities in situ using spatial intensity distribution analysis. Proceedings of the National Academy of Sciences 108:7010–7015.

42. Ward, R. J., J. D. Pediani, K. G. Harikumar, L. J. Miller, and G. Milligan. 2017. Spatial intensity distribution analysis quantifies the extent and regulation of homodimerization of the secretin receptor. Biochemical Journal 474:1879–1895.

